# Identification of *Pseudomonas aeruginosa* genetic determinants that connect redox metabolism to alginate biosynthesis

**DOI:** 10.64898/2026.02.05.704140

**Authors:** Soo-Kyoung Kim, Nishad Thamban Chandrika, Ashton Trey Belew, Najib El-Sayed, Sylvie Garneau-Tsodikova, Vincent T. Lee

**Author notes:** Corresponding authors Sylvie Garneau-Tsodikova: Vincent T. Lee.

## Abstract

*Pseudomonas aeruginosa* is a well-known human pathogen that contributes significantly to chronic infections, particularly in cystic fibrosis (CF) patients. During this chronic infection, *P. aeruginosa* undergoes a phenotype change, the inactivation of *mucA*, which leads to the production of exopolysaccharide alginate, known as mucoid, a key virulence factor associated with biofilm formation. This mucoid phenotype allows the bacterium to persist in the lungs of CF patients for the duration of their lives. Previously, we identified ebselen oxide (EbO) as an inhibitor that suppresses alginate production in *P. aeruginosa*. In the current study, we synthesized a series of structural analogs based on EbO and ebselen (Eb) and evaluated their ability to inhibit alginate production. These analogs did have similar or lower inhibitory activity than EbO. The mechanism by which EbO inhibits alginate production remains unclear. We employed RNA sequencing analysis of *P. aeruginosa* treated with inhibitors and identified several candidate genes potentially involved in this inhibitory pathway. Interestingly, we observed that a transposon and in-frame deletion mutants of the candidate genes were defective for alginate production. These findings suggest there are additional requirements for optimal alginate production in conditions that mimic the CF lung beyond the *algD-A* operon.

**IMPORTANCE:** When bacteria encounter the correct conditions, they can dedicate their energy toward a specific function to maximize the function. One example is the low calcium response in *Yersinia pestis* in which the bacteria arrest growth when grown at 37 °C in the absence of calcium because it uses all its energy for type III secretion. Another example is production of alginate by *P. aeruginosa* in the lungs of CF patients that can lead to occlusion of the airways. In both cases, the dedicated use of energy toward type III secretion for *Y. pestis* and alginate biosynthesis for *P. aeruginosa* reduces the ability of the bacteria to multiply. In the lab, suppressors can be easily identified that restore bacteria growth. The suppressor mutations are often located in the operons that are up-regulated and thereby prevent the execution of the energetically costly process. While these results indicate these processes are energetically costly, we still do not understand how the bacteria dedicate their energy toward these processes over other cellular processes such as growth. Previously, we identified ebselen oxide (EbO) as an inhibitor that suppresses alginate production in *P. aeruginosa*, but chemical analogues fail to improve the inhibitory activity. We used RNA sequencing analysis of *P. aeruginosa* treated with inhibitors and identified several candidate genes potentially involved in this inhibitory pathway. Interestingly, we observed that a transposon and in-frame deletion mutants of these genes involved in redox reactions were defective for alginate production. These findings suggest there proteins may shunt energy for optimal alginate production in conditions that mimic the CF lung beyond the *algD-A* operon. Results from *P. aeruginosa* alginate production may inform how other bacteria can similarly focus energy toward specific cellular processes.

## INTRODUCTION

Cystic fibrosis (CF) is a genetic disorder caused by mutations in the cystic fibrosis transmembrane conductance regulator (CFTR) gene^1^, which encodes an ion channel that plays a critical role in regulating sodium, potassium, calcium-activated chloride, and bicarbonate transport^2–4^. Mutations in *CFTR*, particularly class I-III mutations, result in a loss of CFTR function, reduced chloride secretion, and increased sodium and water absorption *via* the epithelial sodium channel (ENaC)^2^. Defective CFTR function in secretory epithelial cells reduces the airway surface liquid, producing a dehydrated mucus layer that causes ciliary collapse and impairs mucociliary clearance of trapped microbes^2, 5^ and the accumulation of abnormally thick and sticky mucus in the respiratory tract^6^. Consequently, pathogens are not efficiently removed from the CF lung, leading to prolonged and recurrent polymicrobial infections^7^. Overtime, the bacterial community in CF lung shifts from predominantly composed of *S. aureus* to dominated by *P. aeruginosa*, which colonizes the airway of approximately 40% of CF patients by the age of 25 (CF report, 2023)^8^. Despite being constantly colonized with microbes, CF patients often experience episodic acute pulmonary exacerbation^9^ in which the patients experience symptoms including coughing, sputum, and associated decrease in lung function. Of most importance is that exacerbation often leads to a downward spiral in lung function^10^. Exacerbation has been associated with high density of *P. aeruginosa*^11, 12^. This increase in *P. aeruginosa* can be due to a number of factors including the acquisition of new strains, changes in the host that allow existing bacteria to expand, and/or mutations in the existing strain that permit outgrowth. Antibiotic treatment that reduces the numbers of *P. aeruginosa* also reduces symptoms, suggesting that elevated numbers of *P. aeruginosa* in the airway burden correlates with worse health outcomes. In some patients, the *P. aeruginosa* strain undergoes a mucoid conversion in which it produces copious amounts of a viscous polysaccharide called alginate. Antibiotic therapy often is unable to reduce the numbers of mucoid *P. aeruginosa* and patients progressively lose lung function over time^13^.

Our understanding of the progression of *P. aeruginosa* within CF lungs comes from sequencing studies. Sequencing of strains from different CF patients reveals that most *P. aeruginosa* strains that colonize CF patients are different, indicating that they are acquired from the environment^14^. In contrast, longitudinal sequencing studies of sequential isolates from the same CF patient reveal that these *P. aeruginosa* are genetically related and suggests that *P. aeruginosa* becomes a chronic resident in each patient. These data likely rule out the acquisition of new strains as a cause for exacerbation. Chronic colonization provides time for *P. aeruginosa* to accumulate mutations in the genome. When an inactivating mutation occurs in the *mutS* gene, which encodes DNA repair protein, the subsequent isolates rapidly accumulate additional mutations throughout the genome. One gene that is mutated is the *mucA* gene that encodes the anti-sigma factor to prevent *algU* activation. Inactivation of *mucA* leads to constitutive activation of AlgU that drives the expression of the *algD-A* operon that encodes the alginate biosynthesis and export machinery. Mutation of *mucA* is the basis for mucoid conversion.

One approach to prevent the negative clinical outcomes associated with mucoid *P. aeruginosa* is to inhibit the production of alginate. Previously, we identified ebselen (Eb) and ebselen oxide (EbO) as small molecule inhibitors that inhibited alginate production in *P. aeruginosa*^15^. Eb is a well characterized selenium-containing compound that has been shown to be safe for human use in several clinical trials and is consider an antimicrobial agent against multidrug-resistant pathogens^16,17,18,19,20, 21,22,23^. Treatment with Eb or EbO reduced alginate production in *P. aeruginosa* harboring pMMB-*algU*^15^. In the current study, a series of EbO and Eb analogs were synthesized and evaluated to improve efficacy. As these compounds showed a similar potency for inhibiting alginate, we sought to identify cellular target(s) of EbO/Eb. Using RNA-seq, we show that there are only subtle differences in gene expression upon treatment with EbO or Eb when the bacteria is grown in lysogeny broth (LB). Using synthetic CF medium 2 (SCFM2)^24^, which better mimics the CF lung environment, we show that *P. aeruginosa* can make 3-fold more alginate and the treatment with EbO can reduce alginate production by over 10-fold. Using RNA-seq, we show that several genes are induced by *algU* expression in SCFM2, which are then reversed by EbO/Eb treatment. We generated in-frame deletion mutations in some of these genes and show that they are required for optimal alginate production. Together, we show that optimal alginate production may require other cellular factors in addition to the proteins encoded by the alginate biosynthesis (*algD-algA*) operon.

## RESULTS

### Generation of EbO and Eb analogs

To determine if improved inhibitory activity toward alginate production by *P. aeruginosa* can be achieved, several analogs of Eb and EbO (**Fig. 1A**) were synthesized using a previously reported protocol^25^. A series of analogs were synthesized with aliphatic and aromatic substituents (indicated by the R in **Fig. 1A**) using the reaction protocols reported in synthetic Scheme S1. To expand the scope of structural diversity and to verify the influence of certain modification on potency, derivatives with different alkyl chains as well as Eb and EbO analogs with different substituents on the phenyl ring were explored. These Eb and EbO analogs consisted of monosubstitution with alkyl, alkoxy, and halogens, located at the *ortho*, *meta*, or *para* positions. Additional disubstituted regioisomers were also synthesized. When tested

**Fig. 1.**
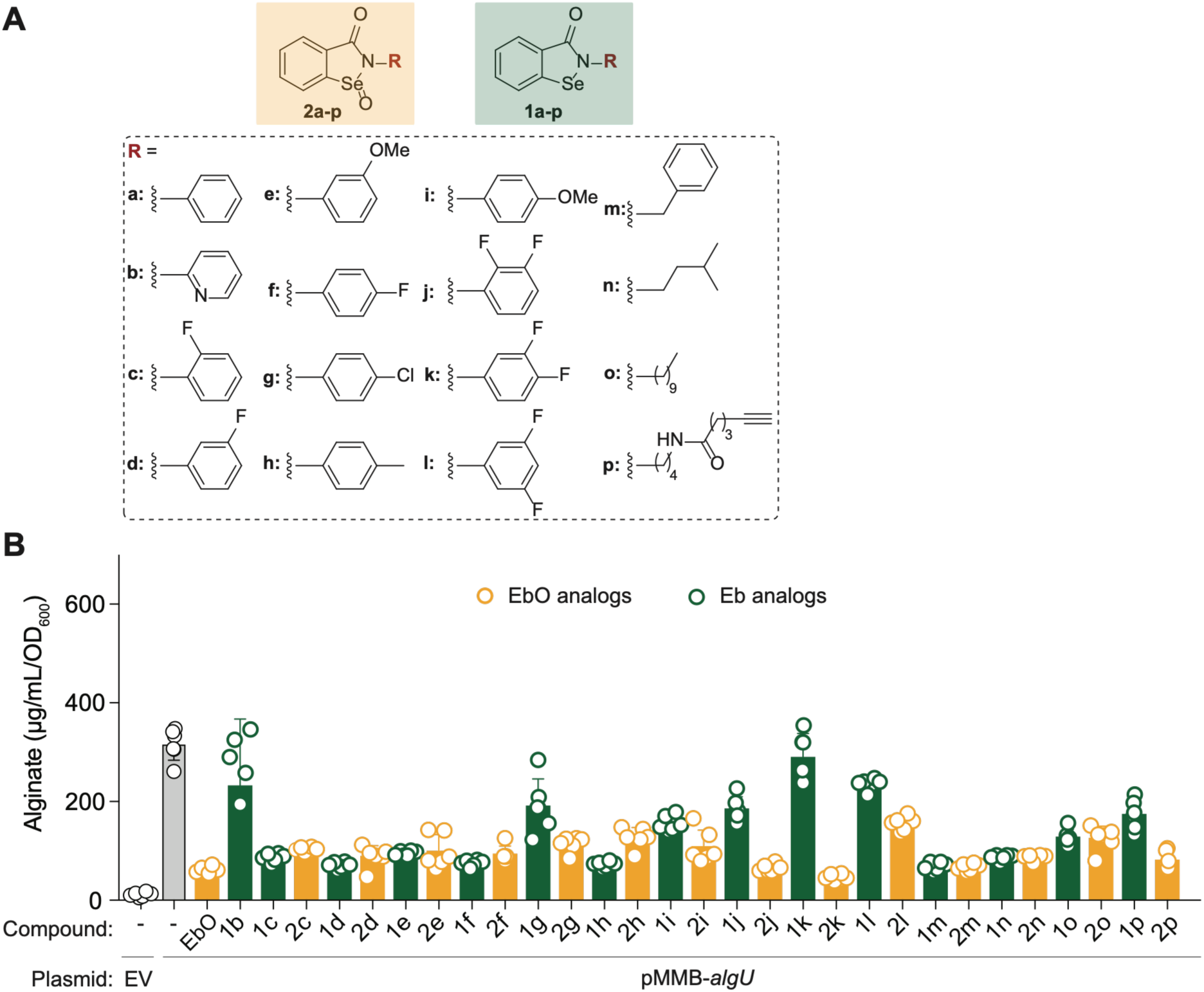
Synthesis and analysis of ebselen (Eb), ebselen oxide (EbO), and their analogs for inhibition of alginate production by *P. aeruginosa* pMMB-*AlgU*. **A.** Chemical structures of compounds used in this study. **B.** Quantification of alginate production in the presence of 100 μM of each of the indicated compound. Graphs show the average of alginate production from 6 independent experiments.

### Testing EbO and Eb analogs for inhibition of alginate production by *P. aeruginosa*

To determine if the analogs synthesized improved inhibitory activity of EbO and Eb, we tested their ability to inhibit alginate production by PA14 pMMB-*algU* induced with 200 μM of IPTG in lysogeny broth (LB – also known as Luria Bertani). In the absence of induction, the bacteria produce less than 10 μg/mL of alginate (**Fig. 1B**, left most bar). Induction of *algU* with IPTG increased alginate production to 316 μg/mL (**Fig. 1B**, grey bar). At 100 μM of EbO, alginate production was reduced by 81% and most chemical analogs had a lower level of inhibition (Fig. 1B). An exception is that compound **2k** demonstrated a slight improvement in alginate inhibition. For Eb analogs, all compounds inhibited alginate production less than EbO.

Compound **2k** and **2j** were tested in SCFM2 (**Fig. S103**) and was shown to be less active than EbO. These results indicated that the Eb/EbO scaffolds are active against *P. aeruginosa* to inhibit alginate production. However, medicinal chemistry has not yet yielded an improved version of the inhibitor. To move towards improved inhibitory activity, we sought to determine the molecular target(s) of alginate production inhibition by EbO.

### RNA-seq of PA14 pMMB-*algU* in LB

To understand the effect of EbO and Eb on PA14 pMMB-*algU*, we extracted total RNA from cells grown in LB with and without IPTG induction and with either DMSO solvent carrier or 100 μM of EbO or Eb. After rRNA depletion, the remaining RNA were made into sequencing library and sequenced by using Illumina HiSeq. Reads were processed for quality control, trimming, and mapped to the PA14 reference genome. The number of reads per gene was tabulated and used for analysis. From this analysis, the IPTG induction increased expression of 112 genes by more than 8-fold and 187 genes were down-regulated by more than 8-fold (**Table S1**). As expected, the *algD-A* operon was induced by IPTG of *algU* (**Fig. 2**, panel 4)^26^. The principal component analysis (PCA) supports that these findings as the main differences in these samples stem from the induction by IPTG rather than treatment with inhibitors (**Fig. 2**, right panel). This *algU* induction was not reversed by the addition of EbO or Eb, suggesting that the reversal of alginate production is due to other processes. Of these differentially regulated genes, we filtered for those genes in which their expression was fully or partially restored to uninduced conditions. For EbO and Eb, while no down-regulated genes recovered expression by 4-fold or more, there were some genes that recovered expression between 2- to 4-fold (**Table S2**). Of the 112 up-regulated gene, there were 4 or 1 gene(s) that increased by treatment with EbO or Eb, respectively. In addition, 17 genes increased transcript abundance by 2- to 4-fold when treated with EbO or Eb (**Table S2**). Of the regions that we identified to be reversed by EbO or Eb, we mapped them onto the circular genome of *P. aeruginosa*. Two regions, *PA14_72650* to *PA14_72690* and *PA14_72710* and *gbuA*, were up-regulated by IPTG and this induction was partially reversed by EbO or Eb treatment (**Fig. 2**, panels 2 and 5). Two regions, *PA14_70650* to *glcD* and *PA14_08000* to *PA14_08300*, were down-regulated by IPTG and reversed by treatment with EbO or Eb (Fig. 2, panels 1 and 3). While these regions were identified, the changes that were observed were restricted to very few genes with smaller magnitude of reversal in gene expression. Thus, we sought to continue our studies using a more physiological relevant medium.

**Fig. 2.**
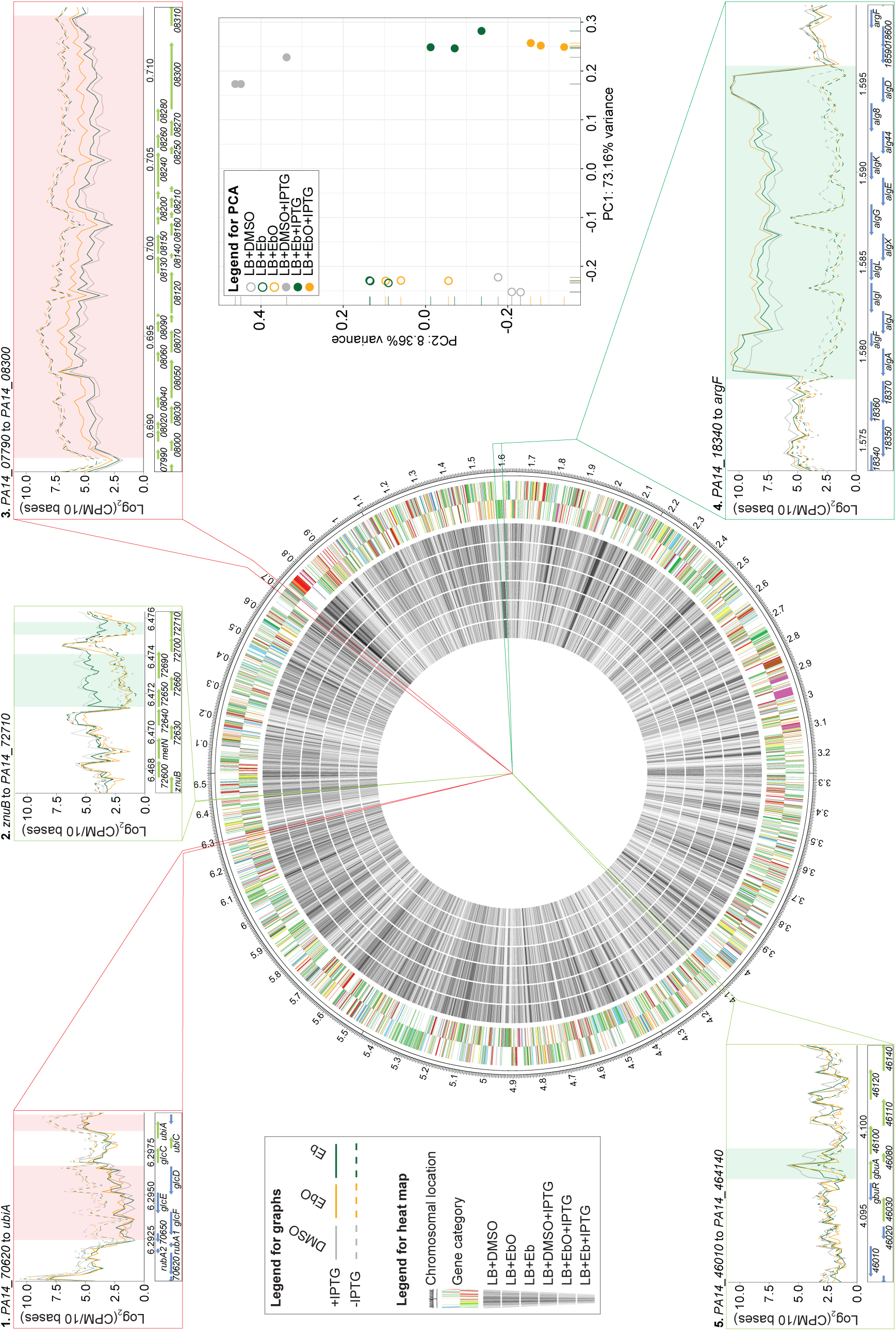
RNA-seq of *P. aeruginosa* pMMB-*algU* grown in LB and treated with EbO or Eb. The RNA-seq data is mapped to the genome and displayed as a heat map. Data for each of the 6 conditions are shown concentric circles with the legend indicated in the Legend for graphs. PCA plot is shown for the triplicate samples of all six conditions. Panels 1-5 show the zoomed in display of log_2_(CPM per 10 bases) at each genomic location for each of the conditions. Segments highlighted in green are induced by IPTG and highlighted in red are repressed by IPTG.

### Inhibition of alginate production in modified synthetic CF medium (SCFM2)

To determine if EbO and Eb can inhibit *P. aeruginosa* in a medium that better mimics the CF lung environment, we tested these inhibitors in modified SCFM2^24^ in the absence of mucin and extracellular DNA as these polymers interfered with alginate assay. PA14 pMMB-*algU* did not produce much alginate in the absence of IPTG induction (**Fig. 3A**). When induced with IPTG, the bacteria produced 49-fold more alginate, which is more than double what we observe in LB (Fig. 3A vs 1B). Treatment with either EbO or Eb reduced alginate production by over 96% as compared to the DMSO control (**Fig. 3A**). Previously, we observed that induction of *algU* with IPTG in LB reduced the number of CFU in the medium as the bacteria convert energy towards alginate production^15^. We tested whether this is also true in modified SCFM2. We found that the CFU numbers decrease when the bacteria were induced with IPTG, in a similarly manner as observed with LB (**Fig. 3B**). Treatment with EbO or Eb restored growth (**Fig. 3B**). Together, these results suggest that EbO and Eb inhibit alginate production in conditions that mimic the CF lung more than what we observed in LB.

**Fig. 3.**
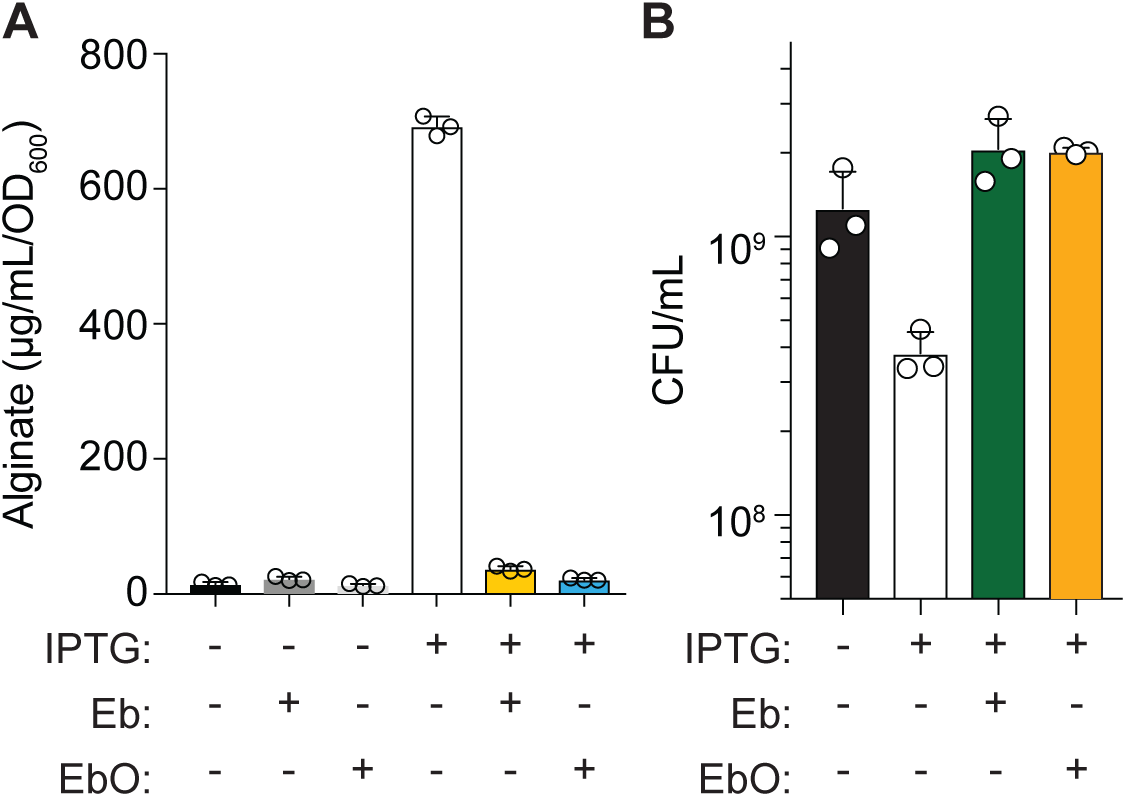
EbO and Eb inhibit alginate production in modified SCFM2 by *P. aeruginosa***. A.** Quantification of alginate inhibition by EbO or Eb with or without IPTG induction of *algU*. **B.** Determination of colony-forming unit (CFU) per mL with or without IPTG induction in the absence or presence of 100 μM of EbO or Eb.

### RNA-seq of PA14 pMMB-*algU* in SCFM2

Because EbO and Eb demonstrated greater inhibitor activity in SCFM2, we performed RNA-seq in SCFM2 for PA14 pMMB-*algU* in the presence and absence of IPTG induction. After tabulating the reads per gene, the number of reads were mapped to genome (**Fig. 4**, chromosome and heat map). IPTG induction led to 467 genes that were up-regulated and 200 genes that were down-regulated (**Table S3**). For the *algD-A* operon, induction with IPTG increase a subset of the genes in the operon by 4-fold, while the remaining genes had similar mRNA levels (**Fig. 4**, panels 5). Addition of EbO and Eb further enhanced expression of the *algD-A* operon suggesting that inhibition occurred outside the expression of these genes. To identify possible targets of inhibition, we filtered for genes that were up- or down-regulated and that verified that this regulation was reversed by treatment with EbO or Eb. Of the 200 down-regulated genes, the expression of only one gene was partially reversed by more than 4-fold by either EbO or Eb, but the gene was not the same one in the two conditions tested (**Table S4**). Of the 467 up-regulated genes, the expression of 21 and 4 genes were repressed by addition of EbO or Eb, respectively. Importantly, all 4 genes repressed by Eb were also found in the EbO treatment (**Table S4**). These IPTG up-regulated genes that were reversed by addition of EbO or Eb are mapped to the genome and expanded in the following panels of Fig. 4: 1. the *cysA* and *sbp* genes, 2. the *nir* and *nor* operons (*PA14_06640* to *PA14_06880*), 3. the *pch* operon (*PA14_09200* to *PA14_09320*), 4. *prpL* gene, and 6. *PA14_36200*. If the effect of EbO and Eb is to alter gene expression, then these 5 sets of genes should be necessary for alginate production in SCFM2.

**Fig. 4.**
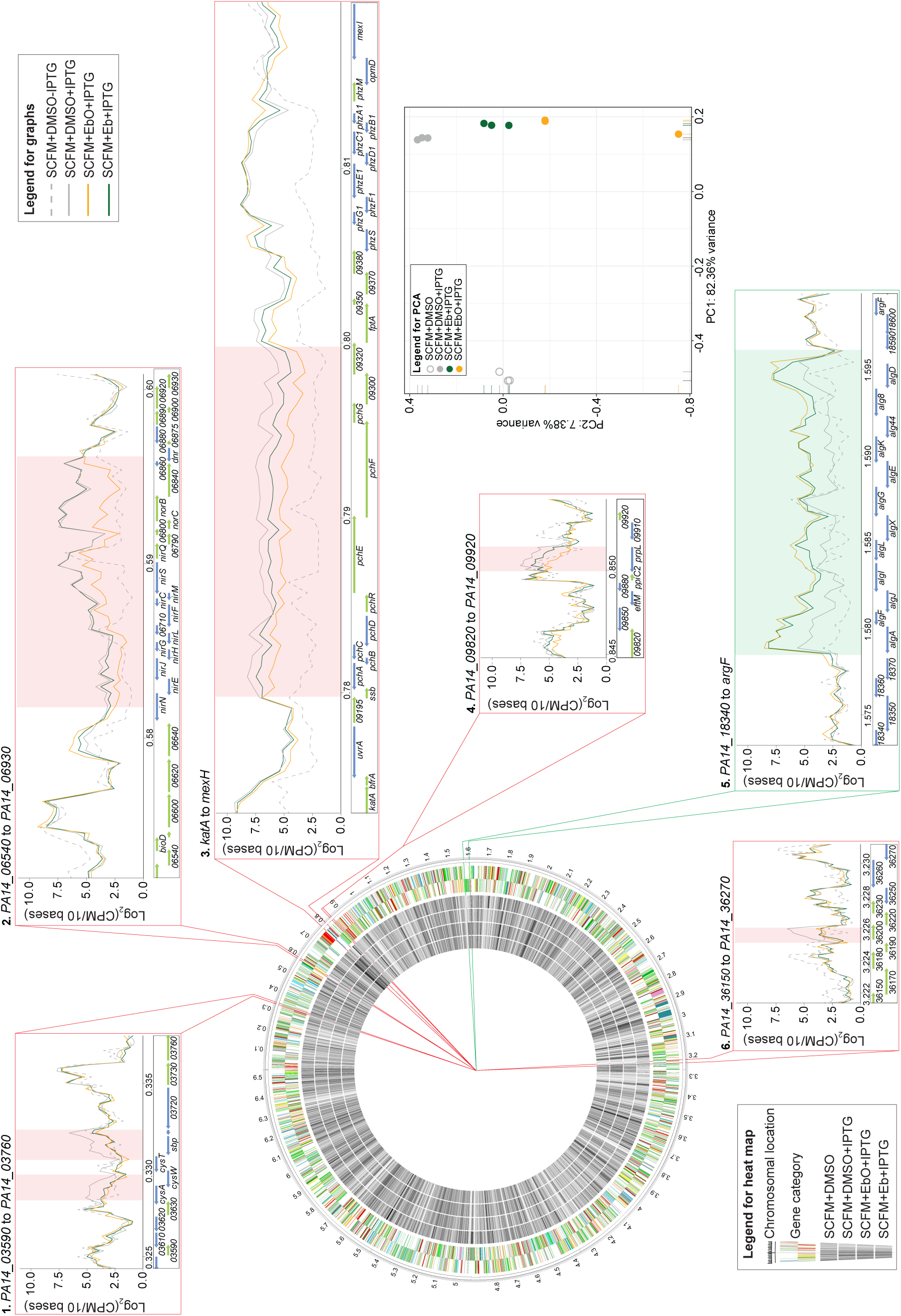
RNA-seq of P*. aeruginosa* pMMB-*algU* grown in modified SCFM2 and treated with EbO or Eb. The RNA-seq data is mapped to the genome and displayed as a heat map. Data for each of the 4 conditions are shown concentric circles with the legend indicated in the Legend for graphs. PCA plot is shown for the triplicate samples of all 4 conditions. Panels 1-6 shown the zoomed in display of log_2_ (CPM per 10 bases) at each genomic location for each of the conditions. The *algD-A* operon is highlighted in green since they are induced by IPTG (panel 5) The remaining genes are highlighted in red since they are repressed by IPTG and reversed by treatment with either EbO or Eb (panels 1-4, and 6).

### Genes identified in RNA-seq in SCFM2 reveal additional requirement for optimal alginate production

To test whether the genes identified from the RNA-seq are required for optimal production of alginate, we utilized the transposon insertion mutants *norB*, *nirS*, *cysA*, *pchA*, *pchB*, *pchC*, and *pchD*. As a control, we used a transposon mutant in *PA14_06600* because it is located near the *nor* and *nir* operons, but its expression is not affected by IPTG nor EbO/Eb inhibition (**Fig. 4**, panel 2). After transforming the pMMB-*algU* plasmid, these strains were assessed for alginate production in SCFM2. Without induction, none of the strains produced alginate. Induction with IPTG led the parental PA14 strain and the *PA14_06600* mutant to make high levels of alginate (>1,000 μg/mL/OD_600_) (**Fig. 5**). In contrast, each of the *norB*, *nirS*, *cyA*, and *pch* mutants produced less than 10% of alginate (**Fig. 5**). These results suggest that the *nor*, *nir*, *cys*, and *pch* operons are required for optimal alginate production in conditions that mimic the CF lung.

**Fig. 5.**
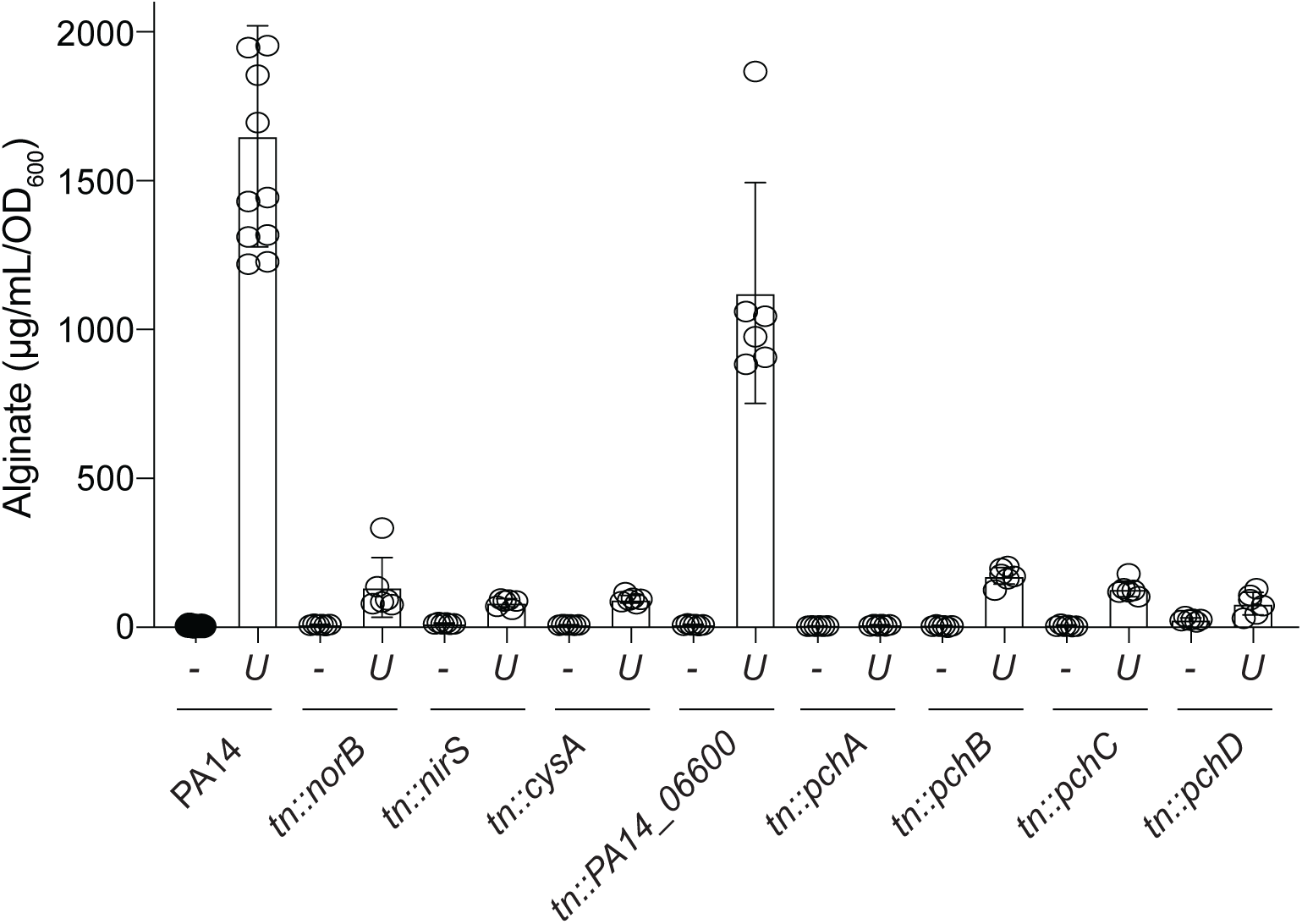
Requirement of the genes identified in RNA-seq for the production of alginate. Transposon mutants were transformed with pMMB or pMMB-*algU* and induced with IPTG for 6 h. Alginate production in was measured in sextuplet and shown with average (bar) and standard error.

### *In vitro* cytotoxicity assay

In order to explore the use of Eb and EbO analogs as alginate inhibitors in CF patients, these compounds were tested for their safety toward mammalian cells. Analogs **1a**, **1d**, **1e**, **1f**, **1h**, **1i**, **1j**, **1m**, **1n**, **1p**, **2c**, **2d**, **2e**, **2f**, **2g**, **2h**, **2i**, **2j**, **2l**, **2o**, and **2p** were tested against A549 human alveolar basal epithelial cells and HEK-293 human embryonic kidney cells to determine their cytotoxic effect at two different concentrations 32 and 64 μg/mL (**Fig. S104**). Against the A549 cells, we observed that none of the analogs displayed toxicity up to 32 μg/mL. At 64 μg/mL, compounds **2c**, **2e**, **2f**, **2g**, **2h**, **2i**, **2j**, **2l**, and **2o** exhibited some toxicity with less than 30% cell survival against A549. In the case of HEK-293 cells, we observed toxicity for compounds **1e**, **2h**, and **2i** at both 32 and 64 μg/mL. Whereas the compounds **1a**, **1f**, **1h**, **1i**, **1j**, **1m**, **1n**, **2c**, **2d**, **2e**, **2f**, **2g**, **2j**, **2l**, and **2o** exhibited some toxicity with less than 30% cell survival at 64 μg/mL.

## DISCUSSION

Despite our previous and current attempts to identify more potent chemical analogs of EbO or Eb, we found that all of the analogs tested had either similar or reduced inhibitory activity toward alginate production (**Fig. 1**). This led us to determine the cellular target that is inhibited by EbO/Eb. RNA-seq analysis of *P. aeruginosa* grown in LB revealed few highly differentially regulated genes that have their gene expression reversed by treatment with EbO or Eb (**Fig. 2**). To better determine if EbO would inhibit in the context of the CF lung, we changed the medium to SCFM2. The PA14 strains induced for *algU* expression produced extraordinary amounts of alginate that was 3-fold higher than in LB (**Fig. 3**). Treatment of EbO or Eb reduced alginate production by over 10-fold suggesting that these compounds are likely functional in the airway of CF patients. RNA-seq analysis of *P. aeruginosa* grown in SCFM2 revealed several candidate genes that had their gene expression profile reversed by the inhibitors (**Fig. 4**). Testing transposon mutants in *norB, nirS, cysA, and pch* mutants showed that these genes are all required for optimal alginate production (**Fig. 5**). Importantly, testing of a transposon mutant in neighboring gene, *PA14_06600*, did not show a similar defect in alginate biosynthesis. Together, this study identified several additional genes needed for optimal alginate production.

This raises a major question regarding the possible contribution of these new proteins for alginate biosynthesis. Alginate is initially produced as a polymannuronic acid from a single precursor monosaccharide mannose-6-phosphate that is converted from fructose-6-phosphate by AlgA^27, 28^. AlgD further oxidize^29, 30^. The remaining genes in the alginate biosynthesis operon are responsible for polymerizing the monosaccharide^31, 32^, transferring the polymer across the inner membrane^31, 33^, modifying the sugar through acetylation and epimerization^34–37^, transport across the periplasm^38^, eliminating polymers that escape into the periplasm^39–41^, and transport across the outer membrane^42 43^. Outside this operon, there is one gene encoding AlgC which is responsible for the conversion of mannose-6-phosphate to mannose-1-phosphate ^44, 45^. Our results presented here are surprising considering that all the functions for alginate biosynthesis from monosaccharide to extracellular polymer have been described.

Our results suggest there may be several additional requirements for alginate production in addition to the alginate biosynthesis operon. To reconcile these two views, we consider the nuanced difference between alginate production and optimal alginate production in the context of the CF lung. This study shows that PA14 expressing *algU* produces alginate production in laboratory conditions such as LB, but the same strain produces more than 3-fold more alginate when grown in SCFM2. This suggests the growth medium can greatly affect the ability of *P. aeruginosa* to produce alginate. Analysis of CF strains with *mucA* mutations have shown a great variation in alginate production^46, 47^ suggesting that there may be additional factors that are strain specific for optimal alginate production.

By characterizing the target of EbO inhibition, we showed that some of these possible factors for optimal alginate production include pyocyanin, pyochelin, nitrate utilization, nitrite utilization, and cysteine production. While these processes are diverse and have no direct chemical connection to the structural components of the alginate polymer, they are all involved in redox processes. Pyochelin is one mechanism in which *P. aeruginosa* acquires iron^48–50^. The *nor* and *nir* genes permit *P. aeruginosa* to utilize nitrate or nitrite, respectively, as terminal electron acceptors ^51^. While it remains unclear what is the function of the *cysAWTsbp* operon in *P. aeruginosa*, but based on homology with *E. coli*, the *cysAWTsbp* genes are required for uptake of sulfate into cells ^52^. Perhaps optimal alginate production requires a drastic shift in funneling the metabolic energy from normal cellular function to the production of alginate. There is evidence that optimal alginate production is costly as we show here and before that there is a near 4- to 8-fold decrease in cell replication. This massive energy shift is well known and often mucoid *P. aeruginosa* isolates propagated in lab that retain mucoid phenotype will accumulate additional mutations to reduce alginate production and thus restoring a growth advantage^53, 54^. Further evidence that there this energy coupling to alginate production exists since Inhibition of alginate by EbO restores growth. This growth restoration does not seem to inhibit the expression of the *algD-A* operon suggesting that the targets lie outside the core alginate biosynthesis genes. Future studies are needed to determine the mechanism by which these additional factors contribute to alginate biosynthesis.

In a push to identify therapeutics to treat various alleles of *cftr*, cell based fluorescent screens for CFTR function utilized a specific variant of Yfp (YFP-H148Q/I521L) that has its fluorescence quenched by iodide ^55^. Expression of this reporter in combination with various alleles of CFTR in cells provided a basis to identify small molecules that restore CFTR function ^56^. These compounds function through two main processes CFTR protein. First, they can correct the shape of the CFTR protein such it can be trafficked to the cell surface ^57^. Second, they can bind CFTR to activate ion transport ^57^. after initial development of treatment for G551D allele, therapeutics for Δ508 was needed as it is the most common allele (ref). High throughput screens identified various compounds and combination of compounds that restored function to Δ508 CFTR ^58^ ^59, 60^. These developments have led to the FDA approved CFTR modulator therapies consisting of either elexacaftor/tezacaftor/ivacaftor (Trikafta) ^61^ or vanzacaftor/tezacaftor/deutivacaftor (Alyftrek) ^62^ that broadly correct the defect in CFTR function for many mutant alleles. Together, these CFTR modulator therapy has been life changing for people with CF by reducing dehydrated mucus in the airways thereby improving lung function. How could EbO/Eb be useful? Despite these successes, *P. aeruginosa* remains in the patients after CFTR modulator therapy ^63, 64^. Whether the chronically colonized microbes including *P. aeruginosa* will reinitiate disease needs to be monitored. As with chronic therapeutics, whether these treatments are effective or begin to lose effectiveness over decades remain to be determined. Lastly, there are specific alleles of CFTR (no protein produced) that cannot be treated with CFTR modulator could also benefit with treatment that reduce alginate production by mucoid *P. aeruginosa*.

## MATERIAL AND METHODS

### Bacterial strains, plasmids, and growth conditions

*P. aeruginosa* strains harboring pMMB-*alguU* were grown in lysogeny broth (LB, often referred to as Luria Bertani broth) containing 50 µg/mL carbenicillin. To test alginate production, *P. aeruginosa* was grown in LB or modified SCFM2 medium^24^ lacking DNA or mucin, but supplemented with 50 µg/mL carbenicillin and 200 mM IPTG at 37 °C. The *P. aeruginosa* PA14 *pchA::tn,* PA14 *pchB::tn,* PA14*pchC::tn,* and PA14*pchD::tn* strains were grown on LB agar plate containing 15 µg/mL gentamicin, then inoculated into LB supplemented with mg/mL gentamicin at 37 °C with shaking overnight. Each transposon mutant was mated with pMMB or pMMB-*algU* on LB agar plates containing 75 µg/mL gentamicin,150 µg/mL carbenicillin, and 25 µg/mL irgasan. The list of strains, plasmids, and primers used in this study are shown in **Table S5**.

### Quantification of alginate production by *P. aeruginosa*

*P. aeruginosa* PA14 harboring pMMB-*algU* were grown overnight in LB broth, sub-cultured with 50 µg/mL carbenicillin and 200 mM IPTG for 6 h at 37 °C in LB or SCFM2^24^. Induced bacteria were collected by centrifuge, the supernatant containing alginate was precipitated by the addition of equal volume of isopropanol at -20 °C. After precipitation by centrifuge for 20 min, the alginate pellet is resuspended in distilled water and assay by the addition of 0.1% carbazole in absolute ethanol and borate-sulfuric acid reagent^65^. The reaction was incubated at 55 °C for 30 min. These samples were measured at 530 nm using the SpectraMax M5 plate reader. This is a spectrometric assay in which the uronic acid sugars were modified by carbazole to produce a compound that absorbs at 530 nm. The values were calculated and converted to unit of alginate production by alginate standard. Alginate concentration was normalized by alginate/mL/OD_600_ of bacterial culture.

### RNA-seq

*P. aeruginosa* PA14 harboring pMMB-*algU* was grown overnight in LB broth, sub-cultured in LB or SCFM containing 50 µg/mL carbenicillin and 200 mM IPTG at 37 °C for 6 h in the absence or presence of 100 mM Eb or EbO. All samples were generated in triplicate. Total RNA was isolated using RNAzol (Molecular Research Center, OH, USA). The total RNA was depleted of *P. aeruginosa* ribosomal RNA using the Illumina total RNA library preparation kit with Ribo-Zero Plus and *P. aeruginosa*-specific oPools (IDT Technologies, IA, USA). The libraries were sequenced on Illumina NextSeq 1000 for 50-bp paired read resulting in 3.3 to 5.9 million reads (average 4.4 ± 0.6 million). Reads were trimmed using Trimmomatic^66^ with a 4:15 sliding window and for a minimum read length of 36. Reads were aligned to the PA14 reference (RefSeq GCF 000404265.1) using HISAT2 (v2.2.1)^67^ and gene counts generated using HTSeq^68^. Discordant reads and unpaired reads were not included in the gene counts. Low count filtering, normalization, and differential expression analysis were performed in R (v4.2.2) via DESeq2 (v1.38.3)^69^. See **Tables S6** and **S7** for the counts per million for each gene for bacteria grown in LB or modified SCFM2, respectively. Sequencing data are deposited in the National Library of Medicine BioProject database under BioProject ID PRJNA1348491.

### Quantification and statical analysis

All data of alginate assay were performed in biological triplicates. Statical analyses were performed using Prism 7.

## ACKNOWLEDGMENT

We would like to thank the Cystic Fibrosis Foundation for funding to V.T.L and S.G.T (LEE22G0 and LEE23G0).

